# Genomic Competition for Noise Reduction Shaped Evolutionary Landscape of Mir-4673

**DOI:** 10.1101/788984

**Authors:** Ramin M Farahani, Saba Rezaei-Lotfi, Neil Hunter

## Abstract

The genomic platform that informs evolution of microRNA cascades remains unknown. Here we capitalized on the recent evolutionary trajectory of hominin-specific miRNA-4673 (Dokumcu et al., 2018) encoded in intron 4 of notch-1 to uncover the identity of one such precursor genomic element and the selective forces acting upon it. The miRNA targets genes that regulate Wnt/β-catenin signalling cascade. Primary sequence of the microRNA and its target region in Wnt modulating genes evolved from homologous signatures mapped to homotypic *cis*-clusters recognised by TCF3/4 and TFAP2A/B/C families. Integration of homologous TFAP2A/B/C *cis*-clusters (short range inhibitor of β-catenin (Li and Dashwood, 2004)) into the transcriptional landscape of Wnt cascade genes can reduce noise in gene expression (Blake et al., 2003). Probabilistic adoption of miRNA secondary structure by one such *cis*-signature in notch-1 reflected selection for superhelical curvature symmetry of precursor DNA to localize a nucleosome that overlapped the latter *cis*-cluster. By replicating the *cis*-cluster signature, non-random interactions of the miRNA with key Wnt modulator genes expanded the transcriptional noise buffering capacity via a coherent feed-forward loop mechanism (Hornstein and Shomron, 2006). In consequence, an autonomous transcriptional noise dampener (the *cis*-cluster/nucleosome) evolved into a post-transcriptional one (the miRNA). The findings suggest a latent potential for remodelling of transcriptional landscape by miRNAs that capitalize on non-random distribution of genomic *cis*-signatures.

## Introduction

MicroRNAs stabilize the evolutionary interface of genotype and phenotype by canalisation of development (Hornstein and Shomron, 2006; Waddington, 1959) and improving the heritability of novel traits (Peterson et al., 2009). Buffering of genetic noise (Raser and O’Shea, 2005) is a central mechanism to improve the robustness of developmental cascades and invoke phenotypic stability by canalisation (Peterson et al., 2009). To regulate genetic noise, miRNAs hybridize to transcripts that encode complementary “seed” sequences and uncouple the subsequent translation of the mRNAs or trigger the endonucleolytic cleavage of targeted transcripts (Brodersen and Voinnet, 2009). As such, genic distribution of the “seed” sequences is a major factor that determines the potential interactions of miRNAs and the system-level manifestations of the interactions.

Random distribution of the seed region in various genes (1 in 4^6^ bp for 6-mer seed regions) and a large-scale mutational drift subsequent to evolution of an individual miRNA can potentially adjust the specificity of the putative interactome to the estimated ≈100 target sites (Brennecke et al., 2005) per miRNA. The latter view, however, conflicts with several major lines of evidence. Unlike 3′-untranslated regions (3′-UTR), sequences in the coding regions (Brummer and Hausser, 2014) are highly conserved and not easily removed by mutational drift from the potential interactome of a newly evolved miRNA. Further, small genes such as those encoding olfactory receptors are not negatively biased against microRNA regulation (Choi et al., 2008) despite lexical simplicity. Finally, emerging evidence suggests the importance of base pairing beyond the seed region in order to improve the specificity of targeting by miRNAs (Broughton et al., 2016). A plausible explanation for complementarity between miRNA-targets is that homologous regions evolve independently and yet simultaneously and are eventually co-opted by evolving miRNAs. In the current paper, we probe the former two propositions, co-option versus *de novo* evolution, by exploring the evolutionary trajectory of miR-4673 (Dokumcu et al., 2018; Farahani et al., 2019a) that is encoded in notch-1 locus.

Recent work in our laboratory demonstrated that miR-4673 can efficiently reprogram the population dynamics of neoplastic cells (Dokumcu et al., 2018). MiR-4673 overrides oncogenic signals that generate heterogeneity in the neoplastic population. To filter the oncogenic noise, miR-4673 induces autophagy leading to the depletion of proteins that perturb cell cycle. One such protein is β-Catenin (Dokumcu et al., 2018), the downstream mediator of Wnt signalling cascade (MacDonald et al., 2009). Wnt/β‐Catenin pathway codes a developmental switch (Hayward et al., 2008) that integrates system-level signalling inputs into a binary outcome (Rezaei-Lotfi et al., 2019). Resolution of the proliferation/differentiation dichotomy under instruction from Wnt/β-catenin pathway (Hayward et al., 2008) is an example of the binarization activity of the cascade.

It therefore is not surprising that altered binarization threshold of Wnt/β-catenin signalling cascade leads to profound morphological novelties. In the nervous system, for example, enhanced activity of Wnt cascade propels significant expansion of the cerebral cortex (Chenn and Walsh, 2002). As such, fluctuations (noise) in the level of free cytoplasmic β-catenin are buffered by multiple parallel post-translational mechanisms (Huber et al., 2001). Yet, unless the transcriptional noise is controlled, fluctuations in post-translational modulators of β-catenin lead to altered binarization threshold of Wnt/β-catenin signalling cascade. Multiple mechanisms have been reported that regulate the activity of Wnt/β-catenin signalling cascade.

A key developmental pathway that regulates the signalling activity of Wnt/β-catenin cascade is the Notch pathway (Hayward et al., 2008). Notch-1 interacts with and suppresses β-catenin (Kwon et al., 2011). Here, we provide evidence for the evolution of miR-4673 and the interacting homologous regions by co-option of transcriptional noise dampeners in notch-1 (Hayward et al., 2008; Kwon et al., 2011) and other post-translational noise modulators of Wnt/β-catenin signalling cascade. The noise dampeners are based on an oscillatory genetic circuit (Brophy and Voigt, 2014) that can regulate the antagonistic post-translational interaction of Notch1 and Wnt/β-catenin pathways (Kwon et al., 2011). In this genetic circuit, input from Wnt signalling can activate the notch-1 enhancer by promoting the assembly of a TCF3/4-β‐catenin complex (Billin et al., 2000). Activation of Notch-1, in turn, reduces the level of cytoplasmic β-catenin in a negative feedback loop (Kwon et al., 2011) and modulates noise in the Wnt signalling cascade. After structural maturation of the notch-1 enhancer into the mature pre-miRNA, a cluster of homologous enhancers in other modulators of the β-catenin pathways were co-opted as its interactome.

## Materials and Methods

### Nucleosome positioning signature identification

An R script was written to calculate nucleosome positioning sequences of genomic sequences based on flexible GG/CC/GC-rich sequences and inflexible AA/TT-rich regions (Valouev et al., 2011). In order to construct the diagram shown in Figure 1a of the main text, changes in NPS-favouring dinucleotide usage frequency in consecutive 40mers of miR-4673 intronic precursor (5ʹ-[A]_4_TGA[T]_2_C[T]_3_…[A]_2_CTGTGAGG[A]_4_TAT-3ʹ] in Notch-1) were plotted according to the formula:

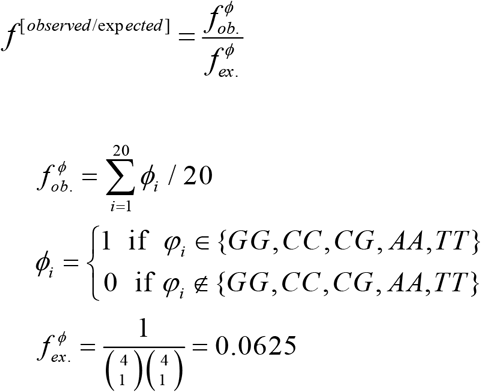

f^φ^_ex._ is the expected frequency of each dinucleotide and *i* is the dinucleotide position in each 40-mer (i_max_=20). The top line in Figure 1b represents ∑(f^φ^_ob._/ f^φ^_ex._)|φ∈(GG,CC,CG) and the bottom line is ∑(f^φ^_ob._/ f^φ^_ex._)|φ∈(AA,TT).

**Figure 1.**
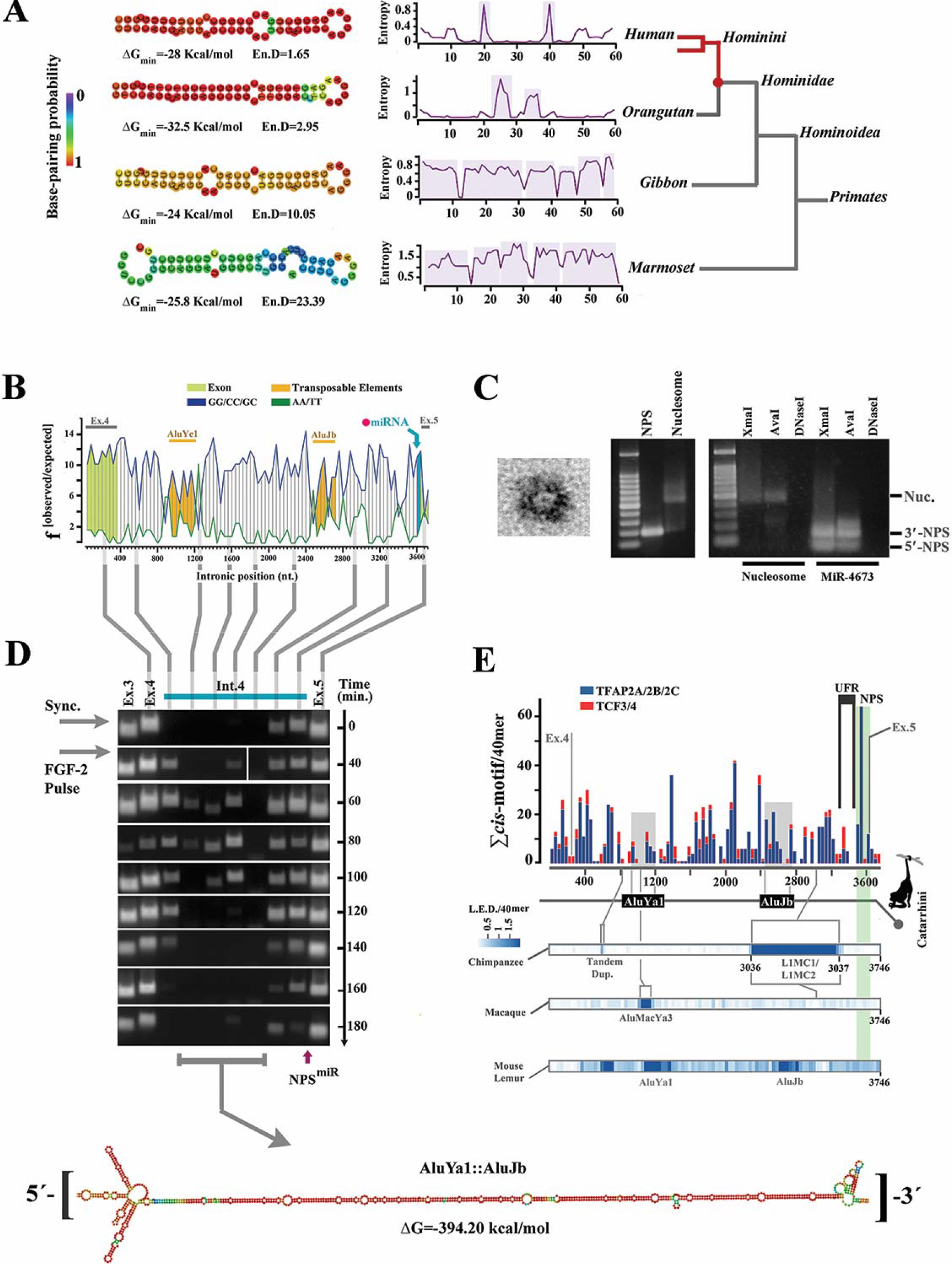
Intronic precursor of miR-4673 in notch-1 codes an active enhancer. **a,** Structural analysis shows improved thermodynamic stability of the RNA hairpin in primate lineage culminating in the structural maturation of miR-4673 hairpin in *Hominins*. **b,** Nucleosome-favouring dinucleotide usage map of human notch-1 intron 4. **c,** Transmission electron micrograph of a nucleosome formed by NPS^miR^ (left). Gel retardation (middle) and restriction enzyme digestion assays (right) provided further confirmation for nucleosome formation by NPS^miR^. **d**, The temporal profile of enhancer-RNAs (eRNAs) that originates from Notch-1 intron-4 (full blots are provided in supplementary file 10). Vertical lines show the location of eRNAs with reference to the dinucleotide usage map. In order to access the eRNA profile, cells were synchronised at G0 and released into G1 by application of a single pulse of FGF-2. Note the temporal stability of 3′-eRNA from a region that corresponds to NPS^miR^ (see supplementary file 11). This region is positioned at 3′-terminus of a stable RNA palindrome formed by two Alu elements. **e,** Histogram shows cumulative distribution of TCF3/4 and TFAP2A/B/C *cis*-motifs in the intron 4 of human notch-1. At the bottom, Levenshtein distance (L.E.D) as a measure of intronic change is aligned to the enhancer map. Sequences that belong to AluJb and AluYa1 insertions in all Simians are in grey. Species-specific transpositional events are marked in LED heat maps.

### SymCurv analysis

SymCurv prediction of nucleosome positioning was performed as detailed elsewhere (Nikolaou et al., 2010; Tilgner et al., 2009). Briefly, curvature values of the sequence were calculated using BENDS(Goodsell and Dickerson, 1994), through a sliding (1bp) window of 30 bp, centered on each nucleotide. From the putative dyads defined as a local curvature minimum, highest SymCurv score, calculated based on the following equations, was selected as the putative dyad of choice:

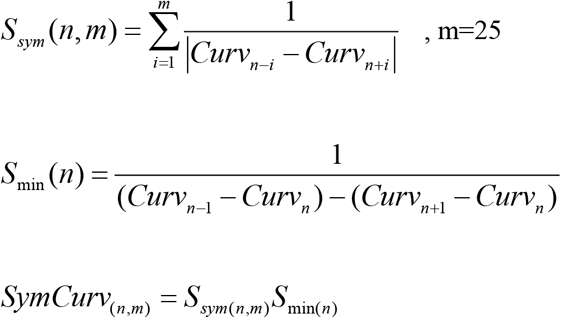

Distance from the putative dyad region (n for SymCurv_(n,m)_=Max) to miR_HR_ was calculated and the two flanking superhelical regions (SHL) accommodating the miR_HR_ were given a score of 1. All other SHL for that particular sequence were given a score of 0. Final score for each SHL was calculated as a sum of individual scores and shown as a radial line graph in Figure 6e.

### Analysis of DNA anisotropy

Analysis of DNA anisotropy was accomplished using nuMap software(Alharbi et al., 2014). The analysis is based on periodic occurrence of AT-containing (WW) and GC-containing dinucleotides (SS) in minor and major grooves and shown to favour rotational nucleosome positioning(Albert et al., 2007; Satchwell et al., 1986; Segal et al., 2006). Briefly, W/S score is calculated as a moving average score centred at position n based on the following equation:

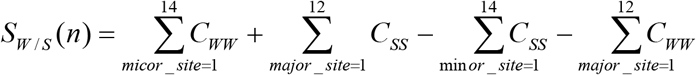

The W/S dinucleotide oscillation plots were generated as sliding window of 1 bp.

### Structural alignment of DNA anisotropy plots

Structural alignment of the W/S dinucleotide oscillations was performed in two steps. Pairwise cross-correlation analysis (see next subheading) of sequences symmetrised to miR_HR_ provided a measure of phase offset which was subsequently curated manually for maximal cross-correlation of oscillations. The distribution of offset values (bps) was presented as a box plot in Figure 6a. The aligned oscillations were subsequently presented as an averaged value (red line, Figure 6a) and bootstrapped confidence interval (grey margin) using ggplot2 library of R platform.

### Cross-correlation analysis

To analyse the cross-correlation of W/S dinucleotide oscillations, structurally aligned W/S oscillations (see previous section) were divided into 20-mers using R platform. Pairwise normalised cross-correlation of the 20-mer was then carried out using the R platform based on the following formula:

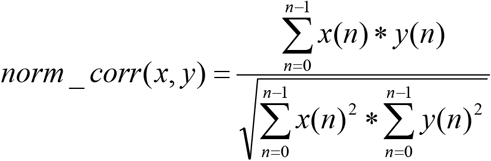

Calculated pair-wise cross-correlation values in each position (each 20-mer) were presented as a linear heat map using ggplot2 library of R.

### PCR amplification of NPS^miR^

Human genomic DNA was isolated from primary human brain progenitors using Qiagen^®^ DNeasy™ Blood & Tissue kit. The DNA fragment corresponding to miR-4673 intronic precursor was amplified using the forward primer TCTTTCAAGCAGGGCGTGTCC and the reverse primer CTCACAGTTCTGGCCGGTGAA and Phusion™ High-Fidelity DNA Polymerase (New England BioLabs^®^). PCR reaction comprised 2 μL of gDNA, 1 μL of 5 μM forward/reverse primers, 4 μL of 5X Phusion HF Buffer and 1 μL of 10 mM dNTPs, DMSO (final concentration: 2%), 0.25 μL of Phusion DNA Polymerase, and 10.25 μL of PCR-grade water.

### Nucleosome reconstitution

Recombinant human histone H2A/H2B dimer and recombinant human histone H3.1/H4 tetramer were purchased from NEB. Nucleosome was assembled with the salt dilution method as described elsewhere(Steger and Workman, 1999). Briefly, recombinant histones (100 pmol of 20μM H2A/H2B dimer, 50 pmol of 10μM H3.1/H4 tetramer) were mixed with the amplified NPS^miR^ (50 pmol, 1:1 octamer to DNA ratio) and adjusted to 2M NaCl and incubated at room temperature for 30 min. The reaction was serially diluted to 1.48, 1, 0.6 and 0.25M NaCl by adding 10 mM Tris EDTA with 30-min incubations at room temperature in each dilution step.

### Gel mobility shift assay

The mobility shift assay was performed as described elsewhere(Ream et al., 2016). Briefly, 10 μl of each sample (reconstituted nucleosome and DNA alone) was mixed with 2 μl of 100% glycerol. A 1% agarose gel was prepared using TB buffer (45 mM Tris, 45 mM boric acid). The agarose gel was run at 10 V/cm for 30 min at room temperature.

### Restriction enzyme protection assay

A restriction map of the NPS^miR^ was prepared using NEB cutter. Two single cutter restriction enzymes (XmaI and AvaI from NEB) and DNAse-I (Life technologies, AM2222) were used to assess the protection of NPS^miR^ by histone in reconstituted nucleosome. The enzymatic digestion was accomplished in a solution comprised of 4 μl of reconstituted nucleosome, 1 μl of restriction enzyme, 1 μl of CutSmart^®^ Buffer and 4 μl of Milli-Q water for 45 min. at 37°C. Digestion with DNAase-I was carried out in a solution comprised of 4 μl of reconstituted nucleosome, 1 μL DNase I (2 U), 1 μL DNase I Buffer, 4 μl of Milli-Q water for 30 min. at 37°C.

### Electron microscopy

Ultrastructural validation of nucleosome reconstitution was achieved by using transmission electron microscopy (Dubochet et al., 1971). Special grids were prepared by carbon evaporation onto a collodion film supported by a carbon film. The solution containing reconstituted nucleosomes was applied to the positively-charged collodion-carbon coated grids for 3 minutes and then stained with 2% uranyl acetate (in water), rinsed 3-4 times in milli-Q water, and dried on filter paper. The grids were then shadowed with platinum from 2 perpendicular directions under an angle of 7-10°. Samples were visualized using a Philips CM120 BioTWIN electron microscope.

### Cell culture

Human neural progenitors (Farahani et al., 2019b) were purchased from ScienCell^®^ (Carlsbad, CA). DMEM/F12 supplemented with 10% fetal calf serum (FCS), recombinant human FGF-2 (20 ng/ml, R&D Systems) and Antibiotic-Antimycotic^®^ (100X, Life Technologies) was used for culturing the cells.

### Enhancer RNA fingerprinting

Several specific short primers were designed to fingerprint the enhancer activity based on the nascent RNA as per Supplementary file 1 and purchased from IDTDNA^®^. The primers avoided two Alu transposable elements in the intron 4 of notch-1 to achieve maximum stringency and specificity. Amplicons were analysed by mfold platform (Zuker, 2003) to confirm lack of significant secondary structure. Post-synchronisation at G0, cells were pulsed with FGF-2 (20 ng/ml) in fresh media and RNA was isolated to fingerprint the enhancer (Klur et al., 2004).

### Reverse transcription and PCR

After DNase treatment, reverse transcription of extracted RNA was carried out using SuperScript-III reverse transcriptase and T4 gene 32 protein (Villalva et al., 2001) (Roche). PCR reactions (35 cycles) were then performed using the primers listed in Supplementary file 3.

### *Cis*-motif profiling

PWMs from JASPAR 2016(Mathelier et al., 2016) were used to identify the TFAP2A/B/C and TCF3/4 *cis*-motifs. A custom R script based on matchPWM function of Biostrings library was written to extract the *cis*-motifs of interest (homology threshold=80%). A histogram was next generated to summarize the frequency of *cis*-motifs. Resolution of the histogram was optimized by adjusting bin width based on the method proposed by Shimazaki and Shinomoto(Shimazaki and Shinomoto, 2007).

### Identification of transposable elements

Two platforms were used to identify and cross-check primate-specific transposable elements in sequences of interest; RAPBASE (Genetic Information Research Institute) and Dfam 2.0(Hubley et al., 2016).

### Palindromic symmetry

CoFold platform(Proctor and Meyer, 2013) was utilized to simulate co-transcriptional folding of sequences of interest. In this platform, the global algorithm of choice for simulating RNA secondary structure was based on thermodynamic parameters proposed by Turner et al(Mathews et al., 1999).

### Karyogram

Circos platform(Krzywinski et al., 2009) was used to generate the circular diagram of Figure 4 of the main text. Isochore map corresponding to each chromosome was also generated using Circos platform by importing the results from GC-profile platform as explained in the following section.

### Isochore mapping of human chromosomes

GC composition of chromosomal segments was calculated using GC-profile platform(Gao and Zhang, 2006). For segmentation of DNA(Zhang et al., 2005) the halting parameter (t0) of 100 and minimum length of 3000 were selected. The output files were subsequently imported into Circos platform to generate the circular isochore maps of Figure 4 in the main text.

### Identification ofmiR_HR_ signature

To identify the miR_HR_ signature, pre-miR-4673 sequence was blasted against the reference library (Homo sapiens all assemblies [GCF_000001405.33 GCF_000306695.2] chromosomes plus unplaced and unlocalized scaffolds in Annotation Release 108). The results were filtered based on defined homology criteria of sequence homology to mir-4673 (Homology_min_>68%, no gaps allowed), conserved [5ʹ-GGCTCCTG-3ʹ] consensus sequence in +/− strand and [miR4673-miR_HR_]^ΔG^<−36 kcal/mol. [miR4673- miR_HR_]^ΔG^ was determined using RNAhybrid platform(Kruger and Rehmsmeier, 2006) (BiBiServ2, Bielefeld University Bioinformatics Server).

### Cytoscape for functional analysis

Cytoscape 3.3.0 was used for gene ontology and functional analysis of miR-4673 interactome genes. ClueGo application was utilised to analyse the identified interactome genes (Figure 4 in the main text, inner circle).

### Analysis of shadow *gga*-Interactome

Shadow interactome of *gallus gallus* were identified based on the same algorithms, criteria and stringency detailed for human genome. Go analysis of chicken interactome was performed using String 10.0 database(Jensen et al., 2009), The Gene ontology Consortium Database(Gene Ontology, 2015), KEGG database(Kanehisa et al., 2017) and PANTHER database(Thomas et al., 2003).

### *Ex-ovo* cultivation of chicken embryos

Fertilised Rhode Island Red eggs were obtained from a local hatchery. The method for ex ovo cultivation of chicken embryos is described elsewhere with slight modifications(Auerbach et al., 1974). Briefly, fertilised eggs were incubated at 37°C with constant rotation for three days before explantation. After this period the eggs were transferred to shell-less culture system comprised of transparent plastic cup, clear polyethylene film and rubber bands to fix the film. The eggs were cracked and contents transferred to the shell-less system and covered by a bacterial agar plate. The system was then incubated at a temperature of 37°C and relative humidity of 70%. Experiments were performed on cultivated embryos at stages HH-16 (51-56 hr)(Hamburger and Hamilton, 1992).

### *Ex ovo* electroporation of chorioallantoic membrane

ECM 830 electroporator (Harvard Apparatus^®^) was used to generate square-wave electric pulses. The solution (20 mM HEPES, 135 mM KCl, 2 mM MgCl2, 0.5% Ficoll 400, 2 mM ATP/5 mM glutathione) containing the pGeneClip-miRNA plasmid (20 ng/μl) was loaded on top of the chorioallantoic membrane and gold-plated Genetrodes (3 mm L-Shape, Harvard Apparatus^®^) were placed on either side of the folded membrane. Electroporation was carried out at 62.5 V/cm, 50 msec, 2 pulses at one-second intervals. The electroporated embryos were then incubated for 24h at 37°C before capturing the photograph followed by harvesting the membrane for histological processing.

### *Ex ovo* electroporation of embryos

ECM 830 electroporator (Harvard Apparatus^®^) was used to generate square-wave electric pulses. The head of the embryo was exposed by cutting the chorionic membrane. A solution (20 mM HEPES, 135 mM KCl, 2 mM MgCl2, 0.5% Ficoll 400, 2 mM ATP/5 mM glutathione) containing the pGeneClip-miRNA plasmid (20 ng/μl) was injected into the canal of the neural tube under illumination in a surgical microscope (Leica M320). Platinum Tweezertrodes (5 mm, Harvard Apparatus^®^) were carefully positioned bilaterally around the embryo’s head and 4 mm apart. Parameters for *ex ovo* electroporation of chicken embryos were adapted from Sauka‐Spengler et al^75^. Electroporation was carried out at 62.5 V/cm, 50 msec, 5 pulses at one-second intervals. The electroporated embryos were then incubated for 24h at 37°C before harvesting for histological processing.

### Data availability

The authors declare that all data generated or analyzed during this study are included within the article and its supplementary information files.

## Results

### Structural features of miR-4673 were instructed by an intragenic enhancer of notch-1

Pre-miRNAs are characterised by a secondary structure (imperfect stem loop) that licences processing by dicer (MacRae et al., 2007) and a primary sequence that instructs the specificity of targeting. We began by addressing evolutionary forces that informed secondary structure of the miRNA in *Hominins* (Figure 1a). The primary sequence of miR-4673 intronic precursor in notch-1 had a bendable G/C-rich core and inflexible A/T-rich flanking regions (Figure 1b). This arrangement is consistent with stringent human nucleosome positioning sequences (NPS) that predict well-positioned nucleosomes *in vivo* (Valouev et al., 2011). The propensity of miR-4673 intronic precursor to communicate with histones was validated experimentally. *In vitro*, a synthetic NPS corresponding to miR-4673 and immediate flanking sequences (NPS^miR^: 5΄-[A]_4_TGA[T]_2_C[T]_3_…TGA[G]_2_[A]_4_TAT-3΄) readily reconstituted a stable nucleosome assembly with recombinant human H2A/H2B/H3.1/H4 histone octamers (Figure 1c). It is known that an additional signature of a high affinity NPS is the palindromic nature of NPS primary sequence that generates superhelical curvature symmetry and accommodates 2-fold dyad symmetry of histone octamers (Thastrom et al., 2004). This feature improves the translational stability of nucleosomes. We reasoned that evolution of the pre-miRNA hairpin might in part reflect structural adaptation of the NPS^miR^ for stable translational localization of a nucleosome through palindromic superhelical symmetry. As translational stability is critical for nucleosomes that protect and regulate access to enhancers (He et al., 2010), we investigated the enhancer activity of intron 4.

Using primary human neural progenitor cells, high-resolution nascent RNA-based temporal fingerprinting of enhancer RNA (He et al., 2010) was employed to access active intragenic enhancers (Arner et al., 2015; Kim et al., 2010) in intron 4 of notch-1. An active enhancer was identified in intron 4 of notch-1 (Figure 1d). The most active 3΄-terminus of the enhancer in intron 4 was mapped to the NPS^miR^ and the upstream flanking region of NPS^miR^ (UFR^miR^: 5΄-[C]_2_[TG]_2_[GA]_2_CA…AGT[C]_2_TGT[C]_2_-3΄). The 3΄-enhancer encoded a combinatorial *cis*-module (Rothbacher et al., 2007) characterized by TCF3/4 binding clusters in UFR^miR^ and TFAP2A/B/C binding motifs in NPS^miR^ (Figure 1e, detailed in supplementary file 2). TFAP2A/B/C family members act as short-range repressors of flanking TCF3/4 (adenomatous polyposis coli) (Li and Dashwood, 2004). The NPS^miR^, on the other hand, can position a +1 nucleosome (Jimeno-Gonzalez et al., 2015) (referenced to the enhancer) that stalls RNA polymerase-II (RNAP-II) (Studitsky et al., 2016) and delays transcriptional elongation. Both these features (short-range repressor and +1 positioned nucleosome) can reduce notch-1 transcriptional noise (Blake et al., 2003) by increasing the activation threshold of the enhancer. Corroborating this notion, we observed more stable and consistent activity (Figure 1d) of the 3΄-terminus of the enhancer (corresponding to UFR^miR^ + NPS^miR^) compared to the 5΄-terminus (exon4-intron4 junction) of the enhancer (Figure 1d).

Enrichment of TCF3/4 cis-cluster in the intron 4 of notch-1 occurred prior to divergence of Simians (Figure 2a). Thereafter, selection of structural features that instruct noiseless Wnt-mediated activation of Notch-1 gained significant momentum in the common ancestor of *Hominoidae* (Figure 2b, c). One such feature was significant enrichment of TFAP2A/B/C family members (short-range repressors of β-catenin (Li and Dashwood, 2004)) (Figure 2b). In *Hominins*, another structural change in the region that corresponds to NPS^miR^ was selection for more stringent histone-independent translational positioning of +1 nucleosome through enhanced superhelical curvature symmetry (Nikolaou et al., 2010) (Figure 2c). Notably, mutational selection for superhelical symmetry of NPS^miR^ did not compromise the rotational nucleosome positioning that relies upon periodic W/S dinucleotide oscillations of the anisotropic NPS (Albert et al., 2007) (Figure 2d). We suggest that structural evolution of the miRNA was as a bystander outcome that reflected selection for the improved superhelical symmetry of the NPS^miR^ with a primary lexicon inherited from the associated cis-cluster (Figure 2e). After evolution, however, the miRNA amplified and expanded the local noise buffering capacity of the NPS^miR^ from notch-1 locus to a global scale through post-transcriptional interactions with other key modulators of Wnt cascade (Figure 2e). In the final step, integration of the miRNA into the existing notch-1/β‐catenin axis resulted in reconfiguration of the cascade by inclusion of the existing components in two feed-forward loops (Figure 3). We next probed the putative evolutionary scenarios for selectivity of targeting by miR-4673 (evolution of the interactome).

**Figure 2.**
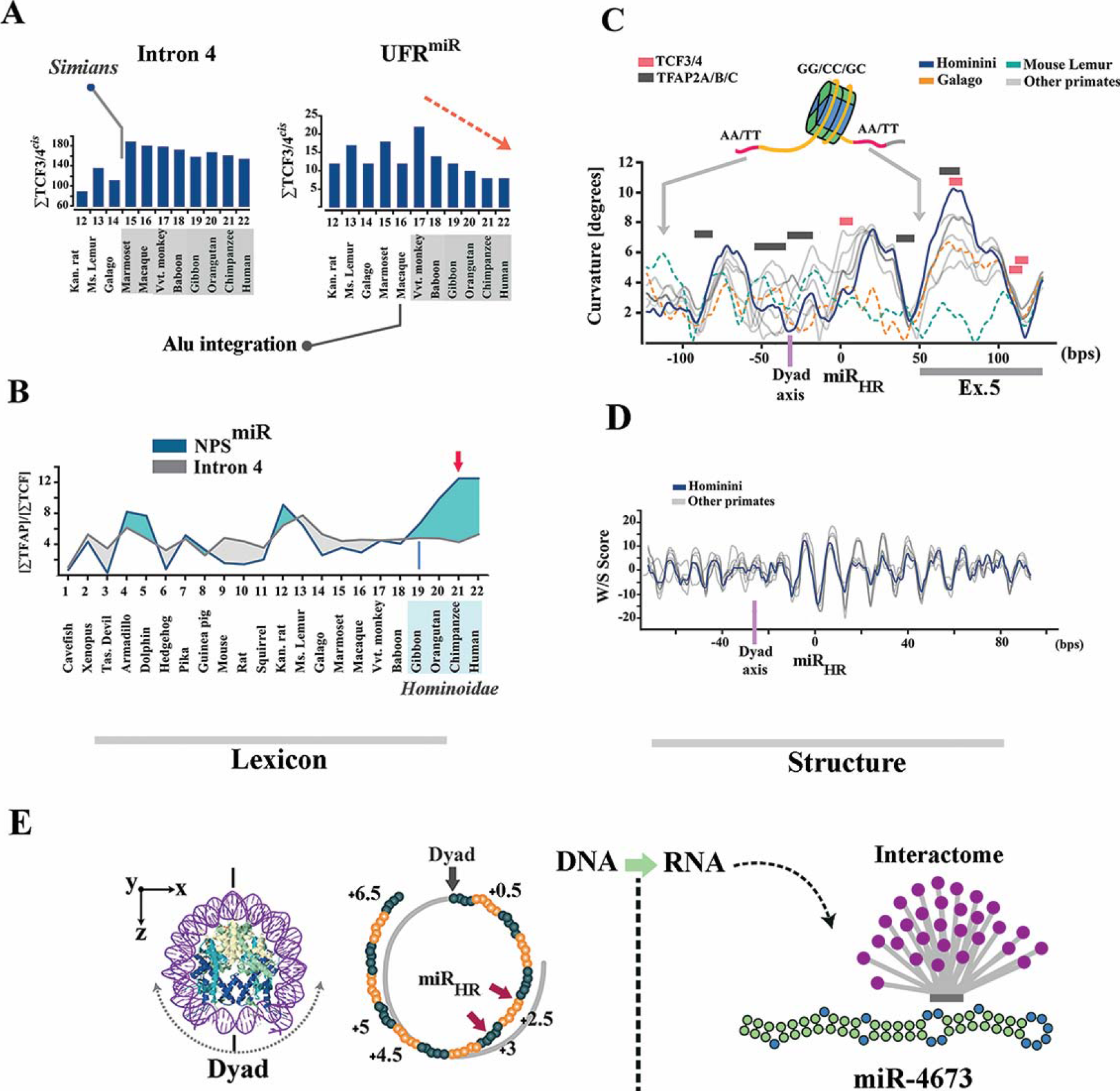
Evolutionary trajectory of the structural features of notch-1 intronic enhancer is aligned to structural maturation of the miR-4673. **a,** TCF3/4 recognition motifs in intron 4 were depleted in higher primates. Depletion of TCF3/4 recognition motifs was more obvious in the upstream flanking region of NPS^miR^ (UFR^miR^) (arrow shows the depletion trend of motifs). **b,** TFAP2A/B/C binding motifs were enriched in NPS^miR^ towards the higher primates. Lines demonstrate [TFAP2A/B/C]:[TCF3/4] ratio in various mammalian species. **c,** In primate lineage, tendency of NPS^miR^ to curve symmetrically gradually increased towards *Hominins* (top diagram, note the increased symmetry of the blue line with reference to the dyad region). This change can improve the translational stability of the nucleosome positioning sequence. **d,** DNA anisotropy evidenced by W/S dinucleotide oscillations did not change significantly in the primate lineage. **e,** The miR_HR_ occupies superhelical locations +2.5 and +3 in the NPS with the palindromic sequence at superhelical locations +4.5 and +5 (middle). The superhelical symmetry accommodates 2-fold dyad symmetry of the nucleosome core particle and associated DNA (left, PDB ID: 3REI). In the transcribed RNA, superhelical positions +1 to +6.5 (65 bp) code pre-miR-4673.

**Figure 3.**
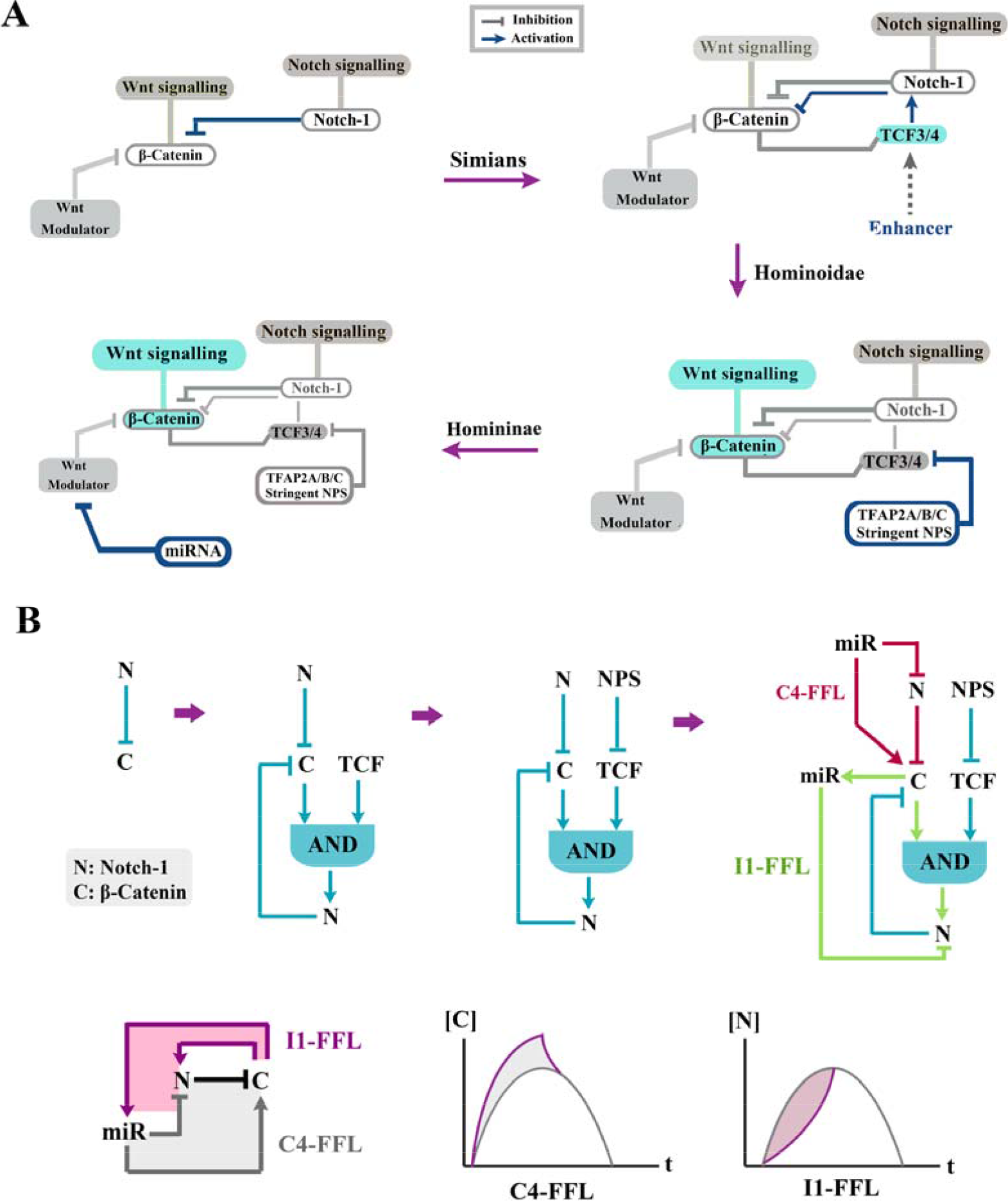
Evolution of network topology of the Notch-1/β-catenin in primate lineage. **a,** Schematic diagrams demonstrate gradual evolution of network topology of the Notch-1/β-catenin axis in prosimians, simians, Hominoidae and Homininae. Note that sophistication of the topology parallels enhances signalling activity of the Wnt/β-catenin in higher primates. **b,** Schematic diagrams (top) demonstrate gradual evolution of the network topology by recruitment of additional network motifs (Alon, 2007; Mangan and Alon, 2003) into the existing circuitry of Notch-1/β-catenin axis. Evolution of the microRNA radically alters (Quarton et al., 2018) the existing genetic network. Evolution of miRNA-4673 in Homininae generates a type-4 coherent feed-forward loop (C4-FFL) that amplifies the activity of the Wnt/β-catenin (bottom). Simultaneously, inclusion of Notch-1 in a type-1 incoherent feed-forward loop (I1-FFL) formed by the miRNA reduces the signalling activity of the notch signalling pathway (bottom).

### Evolutionary trajectory of miR-4673 targets mirrors structural maturation of the miRNA

To resolve evolutionary trajectory of the miRNA targets, we decided to expand the homology requirement beyond the established 6-mer seed region (Broughton et al., 2016). We also expanded the homology region criteria to sequences that are positioned within the intronic, coding and the 5′-untranslated regions (Brummer and Hausser, 2014). The miR-4673 homology regions (miR_HR_) were defined based on sequence homology to miR-4673 (Homology_min_>68%), conserved [5ʹ-GGCTCCTGCC-3ʹ] consensus sequence (+/− strand) and [miR-miR_HR_]^ΔG^<−36 kcal/mol [see methods]. The choice of consensus sequence [5ʹ-GGCTCCTGCC-3ʹ] was based on previous experiments that determined high affinity targets of miR-4673 (Dokumcu et al., 2018). To trace global chromosomal coordinates of putative targets, we used BLAST analysis of miR_HR_ against human genomic sequences (GRCh38). The analysis revealed non-random distribution of miR_HR_ in dense gene clusters associated with GC-rich H3^+^ (GC>52%) isochores (Figure 4). Homology regions were identified in coding sequences (CDS), untranslated exonic regions (5ʹ and 3ʹe-UTRs) and introns (i-UTR) of human miR_HR_-coding genes in sense (miR_HR_^+^) or anti-sense (miR_HR_^-^) strands (Figure 4). Genes signified by miR_HR_ modulated canonical and non-canonical Wnt signalling cascades (Komiya and Habas, 2008) (Figure 5) as detailed in Supplementary file 4. We concluded that miR_HR_ is a signature of a genic cluster whose protein products regulate the cytoplasmic availability of Β-catenin as the main determinant of noise in the Wnt pathway (Figure 5).

**Figure 4.**
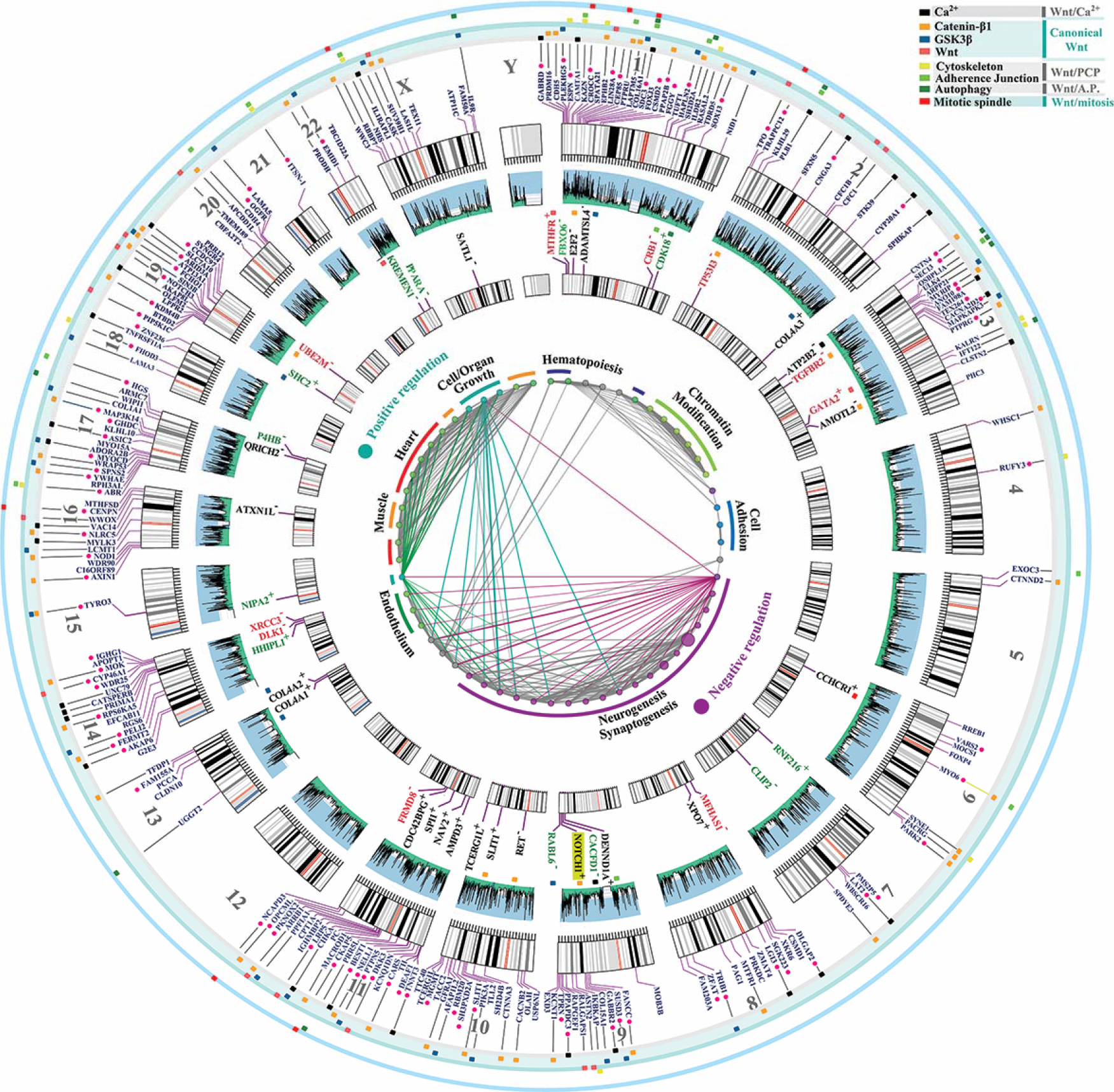
Spatial and functional clustering of homology to miR-4673. Genomic distribution of miR-4673 interactome shows distinct gene clusters localized to H3^+^isochores (middle circular diagram). The miRNA homology regions (miR_HR_) were scattered in coding sequences (inner karyogram genes in black), 5ʹe-UTRs (inner karyogram genes in red), 3ʹe-UTRs (inner karyogram genes in green) and i-UTRs of human genes (outer karyogram). Sense and anti-sense directionality of (miR_HR_) in the inner karyograms is designated as + and -. In the outer karyogram red circles indicate the genes with miR_HR_ in antisense direction to miR-4673. Genes with miR_HR_ signature are involved in regulating development of cardiovascular and nervous systems (inner diagram). These genes act as calibrators (tissue-specific modulators) of Wnt signalling by fine-tuning canonical and non-canonical modules of the cascade (see Supplementary file 4 for detailed elaboration). The outer color-coded layer designates the interactome genes to canonical Wnt and non-canonical Wnt/Ca^2+^, Wnt/PCP (planar cell polarity), Wnt/Autophagy (A.P.) and Wnt/Mitosis cascades.

**Figure 5.**
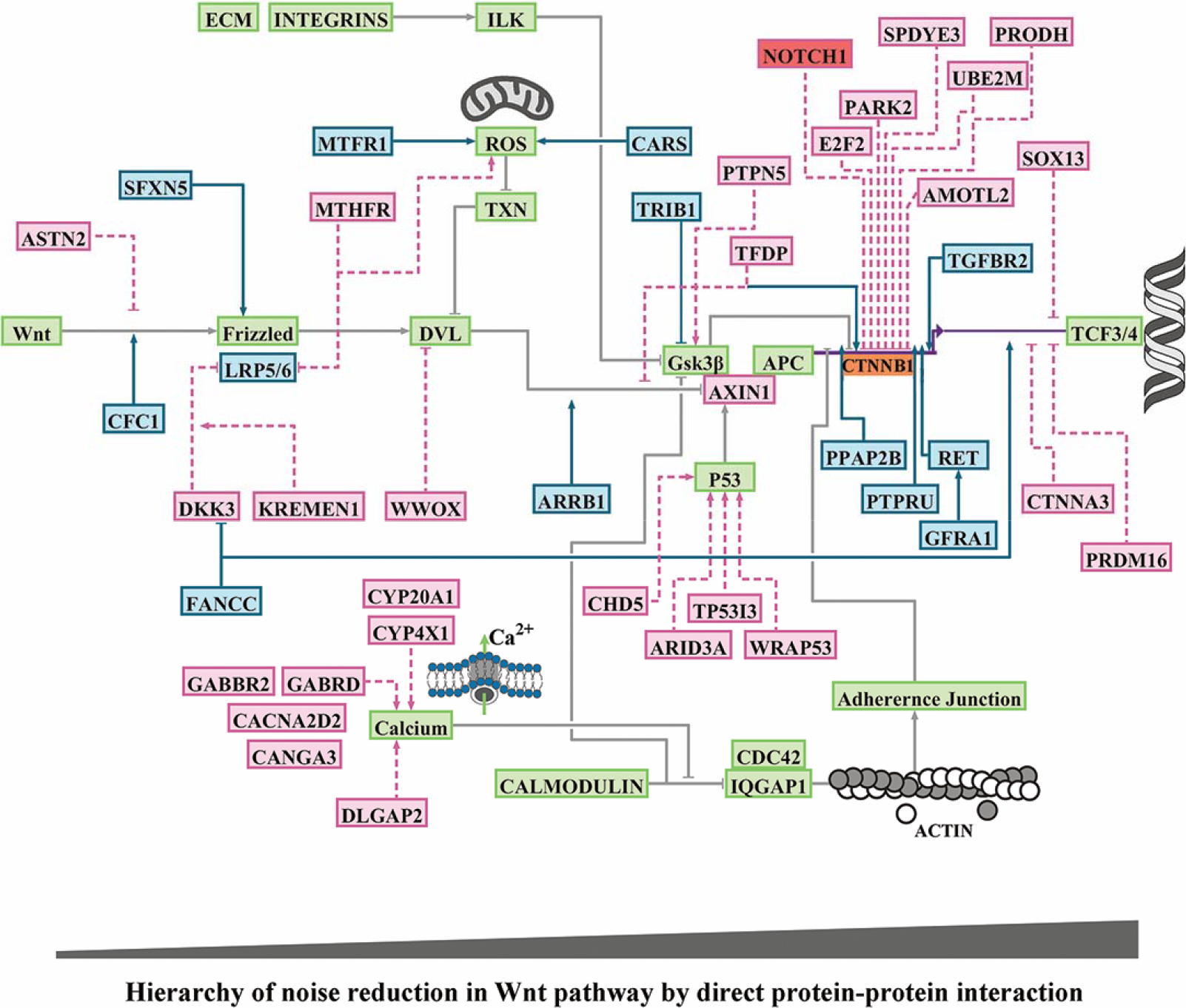
Functional mapping of gene bearing homology to miR-4673. BioTapestry visualization of the Wnt/Β-catenin pathway and the associated modulators. Green boxes demonstrate genes that comprise backbone of Wnt/Β-catenin pathway and that do not code miR_HR_. The blue and red boxes are miR_HR_-coding positive and negative regulators of Β-catenin, respectively (detailed in Supplementary file 4). Genes that code miR_HR_ are mainly involved in direct regulation of Β-catenin. Indirect regulation of Β-catenin occurs by modulating the stability of Cadherin-based junctions that in turn recruit Β-catenin. Another major cluster of genes orchestrate calcium flux and hence the stability of cadherin-based junctions.

To unfold the genomic context of homology regions, all miRNA target regions were symmetrized to miR_HR_ prior to structural analysis (Figure 6a). The aligned sequences demonstrated a G/C-rich central core and inflexible A/T-rich boundary sequences similar to NPS^miR^ (Figure 6a; left). Symmetrisation to miR_HR_ also polarized the aligned sequences into homotypic TCF3/4 *cis*-clusters positioned upstream to miR_HR_ and TFAP2A/B/C *cis*-clusters that flanked miR_HR_ downstream to it in a W/S dinucleotide oscillatory background (Figure 6a). The latter *cis*-anatomy closely replicated that of the NPS^miR^ in notch-1 intron-4 as described previously. We reasoned that targets and the miRNA embarked upon parallel evolutionary trajectories prior to maturation of a functional miRNA. The journey culminated at establishment of bipartite cis-clusters that can coordinate input into Wnt modulator genes by β-catenin (via TCF3/4 module) in a noise-free manner (by TFAP2A/B/C activity and the associated NPS). The miR_HR_ is a chimeric junctional signature of the latter *cis*-modules (Figure 6b); a contention bolstered by symmetrisation of *cis*-clusters to the miR_HR_ (Figure 6a). Notably, genomic regions specialized to recognise TCF3/4 clusters with high affinity can also be recognised by TFAP2A/B/C due to the similarity of the *cis*-lexicon (Figure 6c). The latter observation in part explains diminished randomness in evolution of the described bipartite *cis*-clusters where functional hierarchy in Wnt cascade is aligned to the enrichment of TCF3/4 cis-clusters that in turn facilitates evolution of TFAP2A/B/C cis-clusters. Enrichment of TFAP2A/B/C *cis*-clusters not only can reduce TCF-dependent transcriptional noise but also improves superhelical symmetry of the underlying NPS (Figure 6d, e). Therefore, selection for transcriptional noise buffering by enrichment of TFAP2A/B/C cis-motifs and superhelical symmetry of NPS in the DNA world could empower a parallel and yet dormant journey towards palindromic symmetry in the RNA world (Figure 6e, 7a). We interpreted the parallel structural maturation of the miRNA and the targets as a “competition” for transcriptional noise reduction. After final maturation of miRNA structural signature by notch-1 NPS^miR^, homologous regions in CDS, 5ʹ and 3ʹ e-UTRs of the competing genes were co-opted by the miRNA as its interactome (Figure 4, inner karyogram). To that end, autonomous transcriptional noise filtering capacity of NPS in target genes was amplified and temporally tuned to transcriptional activity of the miRNA and its host gene notch-1. However, conservation of NPS^miR^ in the intronic regions that are not targeted by the miRNA indicates the importance of autonomous transcriptional noise filtering activity of NPS in the target genes.

**Figure 6.**
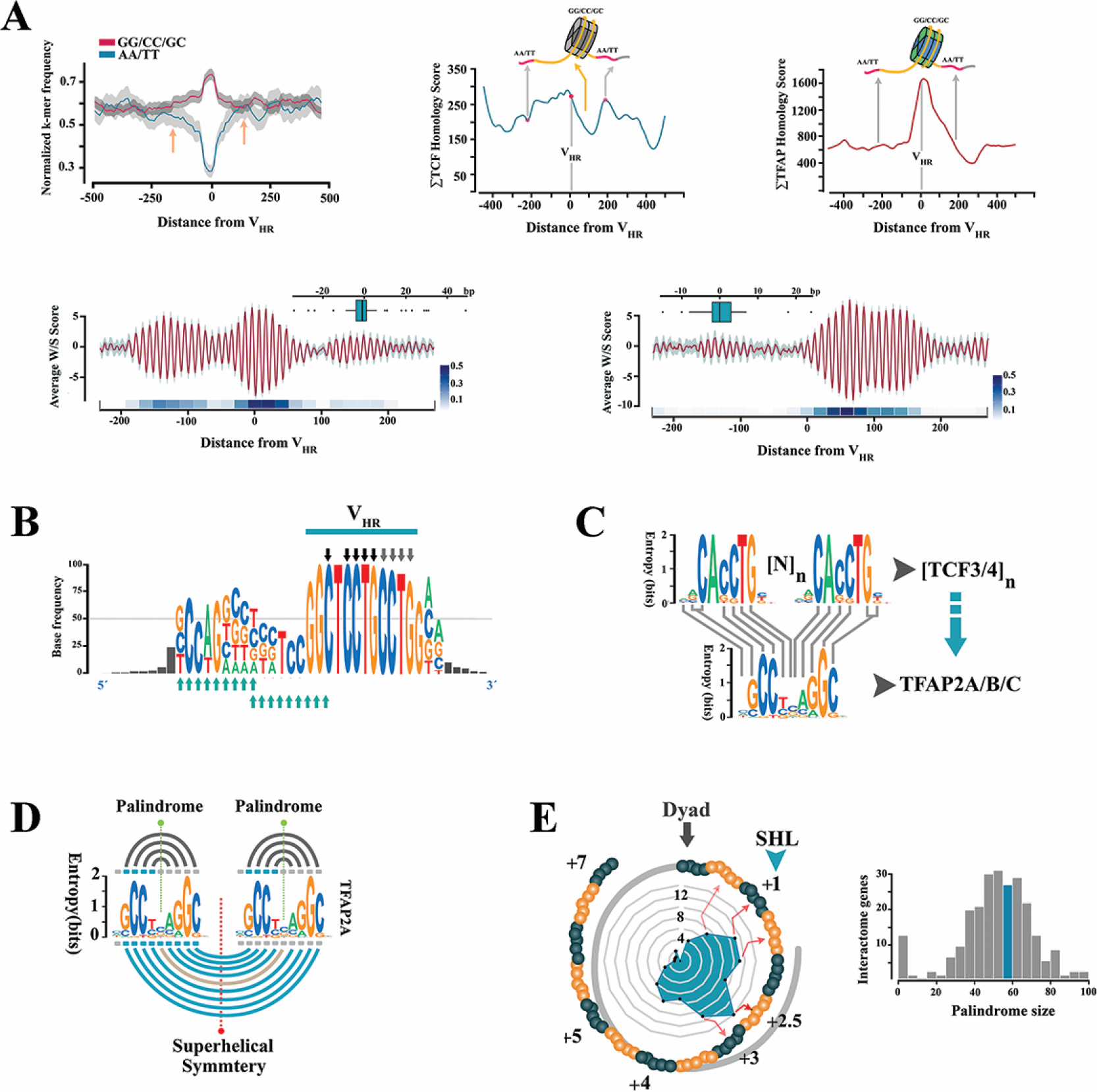
Adaptive evolution of miR_HR_ enhancers in target genes. **a,** Average dinucleotide usage map in target genes after symmetrisation to miR_HR_ (left; arrows indicate AA/TT-rich boundaries). Enrichment of TCF3/4 upstream and TFAP2A/B/C downstream to miR_HR_ in symmetrised sequences (middle, right). Background W/S dinucleotide oscillations (bottom) overlapped the TCF3/4 cis-clusters (left; n=101) or TFAP2A/B/C clusters (right, n=86) (red line: mean, grey margin: bootstrapped confidence interval). Box plots show phase offset between oscillations symmetrised to miR_HR_ (linear heat map; see methods). The linear heat maps demonstrate the average value for pair-wise cross-correlation of 20-mer fragments in structurally aligned NPS sequences. **b,** Consensus motif for the miRNA hybridization region (in RNA world) is chimeric with TCF3/4 cis-motifs (black and grey arrows) in miR_HR_ and TFAP2A/B/C binding motifs (turquoise arrows) upstream to it. **c,** Mutational revision of tandem TCF3/4 *cis*-motifs ([N]_n_: gap between tandem repeats) can generate TFAP2A/B/C recognition motif. **d,** Tandem repeats of palindromic TFAP2A/B/C *cis*-motifs (grey lines) coerce a supersymmetry in the underlying DNA (turquoise lines). **e,** SymCurv analysis of targets with peak TFAP2A/B/C values in the miR_HR_ region (n=46 genes) localized the signature to the superhelical locations (SHL) +2.5 and +3 analogous to the miRNA in Figure 1e (left). The hairpin size that is formed by miR-4673 target regions in the interactome genes (right, Supplementary file 9).

**Figure 7.**
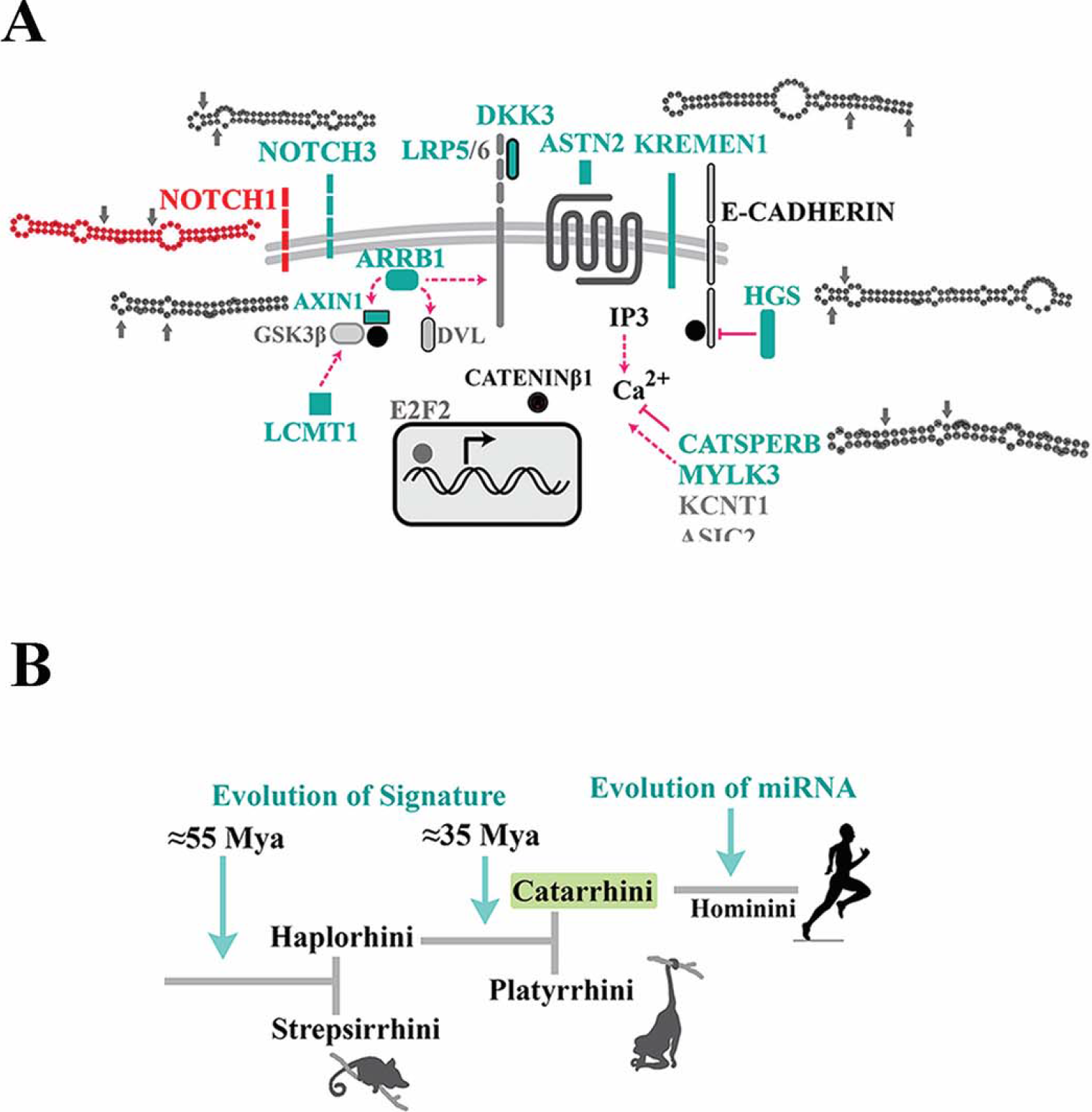
MiR-4673 targets are dormant immature pre-miRNAs. **a,** Near-perfect RNA stem-loops were identified in key modulators of Wnt cascade that are targeted by the miRNA. Selection for structural features of DNA that improve nucleosome positioning propels a parallel journey in the RNA world to superimpose a stem loop signature in the miR_HR_ region of most targets similar to miR-4673 secondary structure. Arrows designate miR_HR_ in the hairpins. **b,** Co-option of transposable elements into *cis*-clusters expanded the extant *cis*-clusters during the Eocene epoch prior to divergence of Catarrhini. Afterwards, a period of dormancy culminated in the evolution of the miRNA.

Reduction of noise in the transcriptional landscape of Wnt modulator genes by recruitment of bipartite enhancers occurred during the Eocene epoch and prior to divergence of Platyrrhine (≈56-33.9 Mya) by mutational remodelling of the extant genomic landscape or integration of transposable elements (Fig 7b, Supplementary file 5). This phase ceased before divergence of AluY transposable elements (Batzer and Deininger, 2002) in Simians (Figure 7b, Supplementary file 5). Thereafter concurrent with the Eocene–Oligocene extinction event (~34 Mya) (Ivany et al., 2000) progressive selection of features that enhance translational stability of nucleosomes, e.g. superhelical symmetry and AA/TT boundary constraints, occurred. To uncover the selective advantage that stabilized miR-4673 in hominins after Pliocene, we investigated post-transcriptional reduction of genetic noise at a high temporal resolution.

### Activity of miR-4673 temporally complements the noise-filtering capacity of notch-1

We inferred the impact of miR-4673 on transcriptional noise based on relative temporal availability of the targeted transcripts with reference to cell cycle in the control and the transfected cells. The transcriptional activity of genes that encoded miR_HR_ in coding sequences and untranslated 5ʹ and 3ʹ-exonic regions (genes of inner karyogram in Figure 4) was measured using real-time qPCR. Preliminary experiments demonstrated relative synchronisation of the population-level cell cycle after 20 hours at G0 phase of cycle (Dokumcu et al., 2018). Based on preliminary experiments, the first measurement (t_0_) was carried out 19 hours after electroporation of the cells with naked miR-4673 (200 nM/10^6^ cells). Thereafter, transcriptional activity was measured at regular intervals every 20 minutes for the next three hours. Transcriptional activity of cyclin-dependent kinase-1b (cdkn1b) and cyclin-D1 long isoform (ccnd-1l) was measured to fingerprint the phases of cell cycle precisely (Figure 8a). Cyclin-D1 is required for progress from G1 to S phase (Ezhevsky et al., 1997) and cdkn1b (p27) regulates the assembly and activation of CDK4 and Cyclin D1 (Larrea et al., 2008). Therefore the transcriptional activity of these two genes increases immediately after M phase to propel progress in G1 phase. We noted a significant boost in the activity of ccnd1l and cdkn1b at t_100_ (5th time point or t=100 min) and t_120_ (t=120 min) (Figure 8a). This transcriptional fingerprint was consistent with population-level transition from M to G1 phase of cell cycle at t_100_. On the other hand, completion of M phase required ≈30 min as measured by high-frequency single cell tracking of proliferating cells (Figure 8b). Hence, the temporal window from t_80_ to t_100_ corresponded to an M phase-rich period in the population of cycling cells. The temporal points prior to t_80_ and after t_100_ were mapped to G2-rich and G1-rich windows, respectively.

**Figure 8.**
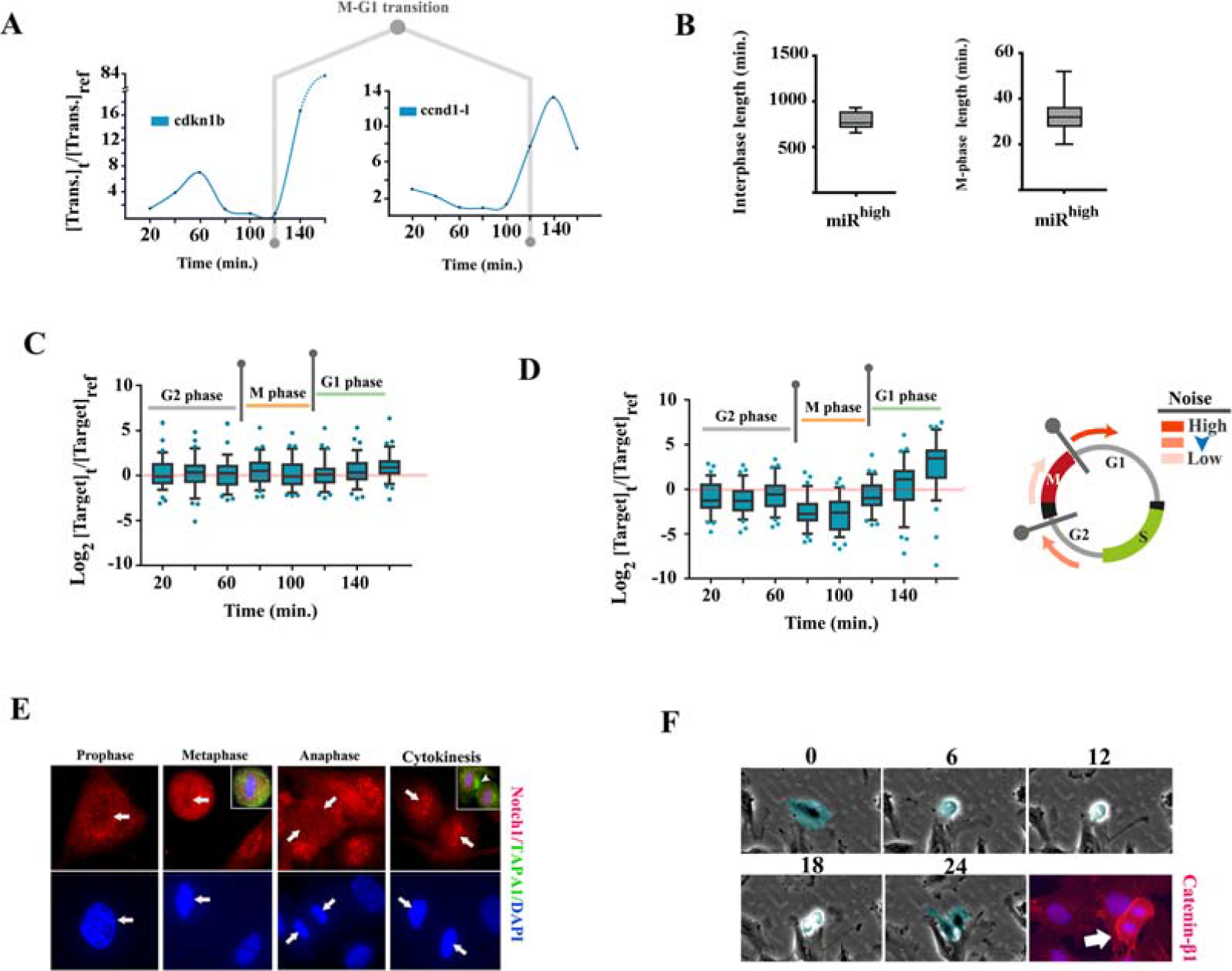
Cell cycle-specific modulation of transcriptional noise by miR-4673. **a,** Transcriptional fingerprint of cyclin-dependent kinase-1b (cdkn1b) and cyclin-D1-long isoform (ccnd1-l) was used as a proxy for cell cycle profile of the dividing population. The rising levels of ccnd1-l transcript at t_120_ is consistent with M-G1 transitional phase. **b,** The length of M phase and interphase was measured using single-cell live imaging analysis of dividing cells (n=20 cells). Based on the average length of M phase, t_80_ and t_100_ temporal windows accommodate the transcriptional profile of M phase. **c,** Each box plot demonstrates the pooled expression levels of miR-4673 targets (n=43 inner karyogram genes of Figure 4) at specific time points ([Target]_t_) normalized to the expression levels of the same genes at time zero ([target]_ref_). Note that in the absence of exogenous miR-4673 the temporal expression profile shows a slight depression at t_100_ followed by an increase at t_160_. **d,** Application of exogenous miR-4673 amplifies the temporal transcriptional signature of miR-4673 targets. Amplified suppression of the transcriptional profile prior to t_120_ anticipates the marked increase in the expression level of targets at t_140_ and t_160_. **e,** Notch-1 fingerprinting shows absence of the activated protein in the nuclear compartment during M phase and the re-emergence concurrent with cytokinesis (top left). Cell surface glycoprotein, TAPA-1, demarcates the contractile ring formed during cytokinesis. **f,** Single cell tracking of proliferating cells demonstrated increased level of cytoplasmic β-catenin after mitotic cell rounding and prior to cytokinesis (numbers: time points in min.).

Following the amplification of miR-4673 (miR^high^ cells), we noted a bimodal reprogramming of the transcriptional landscape of the targets (Figure 8c, d). Amplification of the miRNA suppressed the transcript availability of target genes particularly at t_80_ and t_100_ (Figure 8d, detailed in supplementary file 6). Hence, maximum transcriptional suppression by miR-4673 of the targets occurred during M phase of cell cycle (t_80_ to t_100_). Notably, concurrent with entry into early G1, we noted enhanced transcriptional activity of the targets in the miR^high^ cells (Figure 8d). This finding is consistent with the reported cell cycle-dependent switching of miRNA activity from being a silencer to an activator (Vasudevan et al., 2007). The maximum noise buffering by miR-4673 that occurs during M-G1 transition compensates for the absence of noise reduction by notch-1 (miR-4673 host gene) during mitosis (Figure 8e). Concurrent with mitotic exit, the cytoplasmic level of β-catenin protein suddenly increases (Figure 8f). This is partly due to suppression of β-TrCP1-Skp1 complex during M-G1 transition (Wu et al., 2003) that enhances the level of β-catenin to propel cell cycle progression (Shtutman et al., 1999). If the binary threshold for β-catenin availability is surpassed during the M-G1 transition, the cell commits to the next mitotic cycle by progress into G1. Therefore, noise buffering activity of miR-4673 during this phase could potentially impact upon proliferative capacity of the cycling cells by modulation of the activation threshold of Wnt/β-catenin pathway. We sought experimental proof for this notion by identification of an interactome in other species that encodes the miR_HR_ but is not yet targeted by any known miRNAs. Such a “shadow interactome” would anticipate and become active after structural maturation of a dormant miRNA and may accelerate acquisition of novel adaptive phenotypes at times of environmental crisis (Figure 9a).

**Figure 9.**
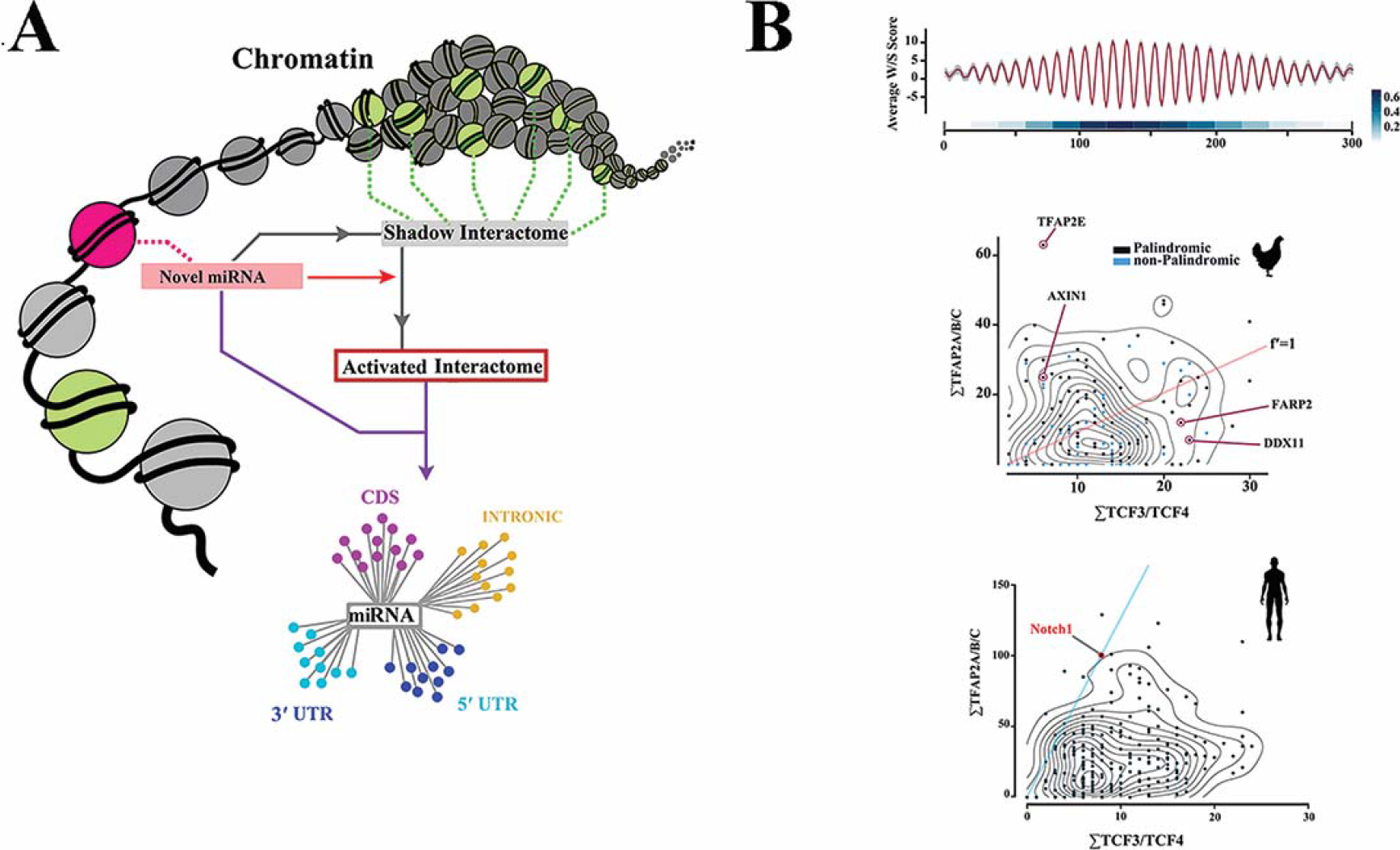
“Shadow interactome” and rewiring of genomic connectivity. **a,** Schematic demonstration of a shadow interactome evolved in a genomic context and in the absence of a functional miRNA. Members of the shadow interactome may complete their structural maturation towards a hairpin that is recognisable by Dicer and co-opt the other members as an activated interactome. b, Genomic features of a “shadow interactome” in *Gallus gallus* with sequence homology to human miR-4673. The miR-4673 homology region in *Gallus gallus* (gga-miR_HR_) showed W/S dinucleotide oscillations characteristic of anisotropic NPS (top; red line: mean, grey margin: bootstrapped confidence interval). The linear heat map demonstrates pair-wise cross-correlation of 20-mer fragments in structurally aligned NPS. Bottom left scatterplot/contour map shows distribution of TFAP2A/B/C and TCF3/4 *cis*-motifs in gga-miR_HR_ gene cluster compared to human miR-4673 interactome. Note that enrichment of TFAP2A/B/C parallels generation of palindromic symmetry (black dots as opposed to blue dots clustered in the vicinity of red line). Also the lower TFAP2/TCF3/4 ratio in gga-miR_HR_ compared to human miR_HR_ suggests less stringent requirement for genetic noise reduction in chicken (red line: equal TFAP2/TCF3/4 ratio).

### Activation of a dormant interactome by miR-4673 reduces developmental noise in chicken embryogenesis

The miR_HR_ was targeted in *Mus musculus* by *mmu*-miR-3104-5p (Supplementary file 7). Hence, *Gallus gallus* (*gga*) was chosen as an accessible, distant endothermic species with distinct GC-rich isochores. An orthologous chicken-specific shadow interactome with miR_HR_ NPS signature embedded in a simple W/S dinucleotide oscillatory background was identified that was not targeted by any known gga-miRNAs (Figure 9b). An interactome map was generated to analyse targets of miR-4673 in chicken (Figure 10 and Supplementary file 8). The gga-specific targets were also identified as modulators of Wnt cascade noise (Figure 10 and Supplementary file 8). *In ovo* electroporation of human miR-4673 into chorio-allantoic membrane (CAM) of chicken embryo (HH16) triggered remarkable expansion of CAM vasculature (Figure 11). Similarly, experimental application of miR-4673 to lateral ventricles of chicken embryo HH 23-25 (E4-4.5) followed by *in ovo* electroporation, triggered extensive synchronised and yet aberrant expansion of neural and hematopoietic precursors (Figure 11). The enhanced mitotic activity was consistent with altered activity of Wnt/Β-catenin pathway that is known to increase proliferative activity during organogenesis (Chenn and Walsh, 2002). The cycle-independent application of the miR-4673 suffers from a lack of temporal fine-tuning due to the constant availability of the miRNA. The experiment, therefore, demonstrates the critical role of microRNA host genes in regulating the temporal dimension of miRNA-target interactions. We concluded that the dormant capacity of a shadow interactome may can be accessed at times of evolutionary crisis by evolving miRNAs to propel remarkable evolutionary adaptations.

**Figure 10.**
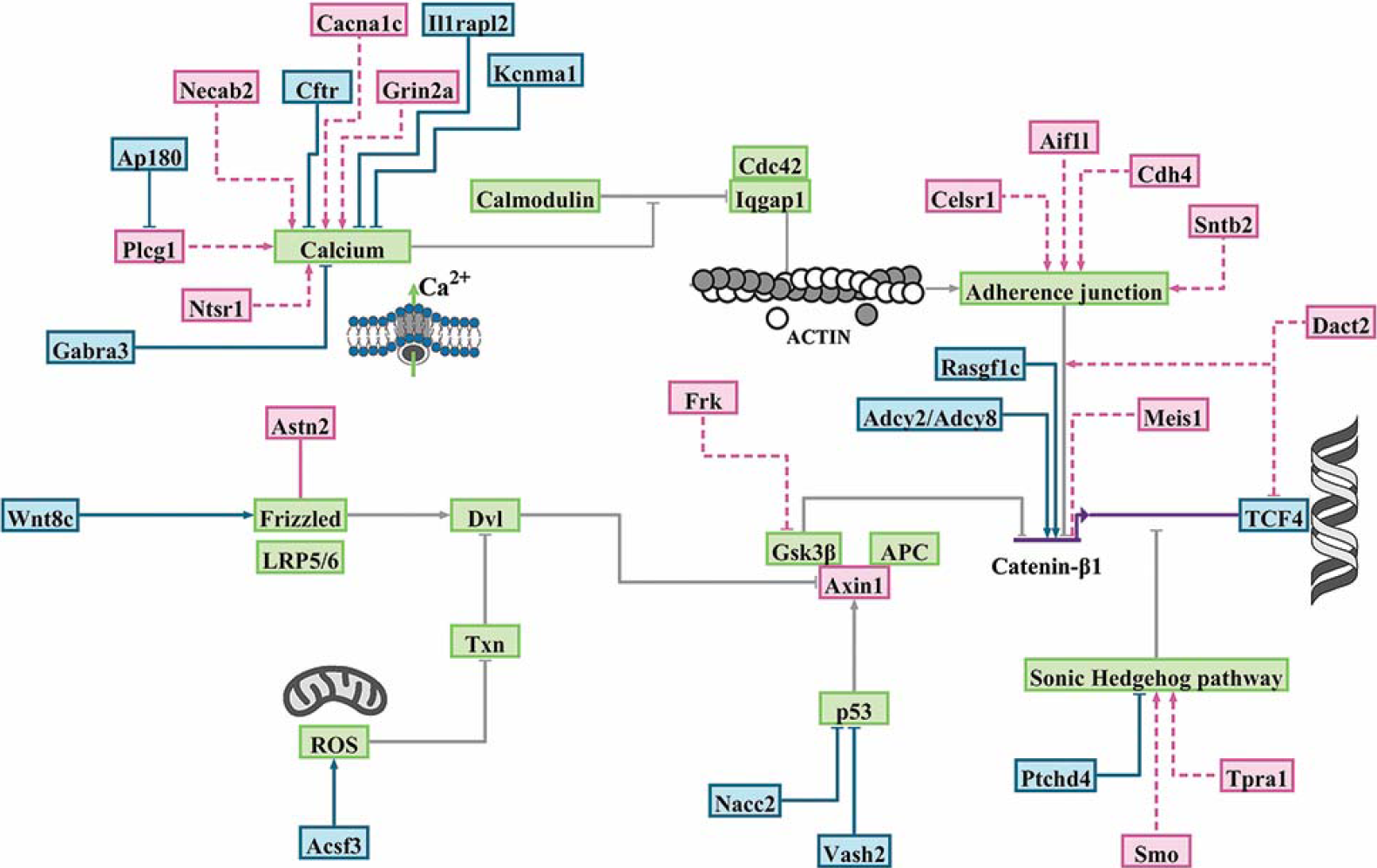
Functional mapping of *Gallus gallus* gene bearing homology to miR-4673. BioTapestry visualization of Wnt/Β-catenin pathway and the associated modulators. Green boxes demonstrate genes that comprise the backbone of the Wnt/Β-catenin pathway and that do not code miR_HR_. The blue and red boxes are miR_HR_-coding positive and negative regulators of Β-catenin, respectively (Detailed is Supplementary file 8). Some of the *gga*-genes that code miR_HR_ are involved in direct regulation of Β-catenin. Most *gga*-interactome genes are indirect regulators of Β-catenin activity. Indirect regulation of Β-catenin occurs by modulating the stability of Cadherin-based junctions that in turn recruit Β-catenin. Another major cluster of genes orchestrate calcium flux and hence the stability of cadherin-based junctions. A small cluster of miR_HR_-bearing *gga*-genes communicate with sonic hedgehog cascade, a major antagonist of Wnt/Β-catenin, to regulate the activity of the latter pathway.

**Figure 11.**
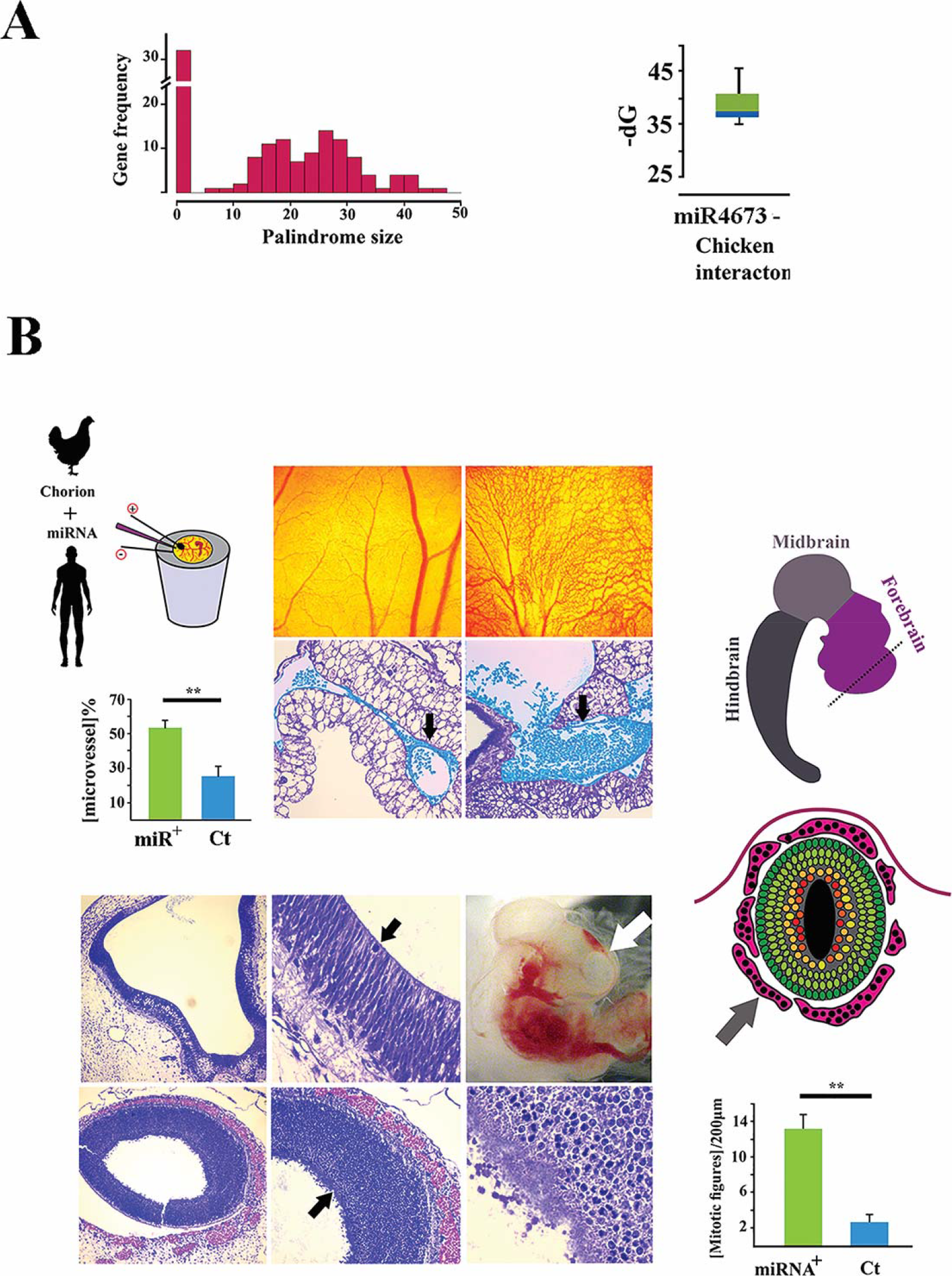
Experimental activation of “shadow interactome” in *Gallus gallus.* **a**, Histogram (top left) shows the distribution of palindrome size in gga-miR_HR_ region. Box plot (top right) shows ΔG for miR-4673/*gga*-miR_HR_ hybridization. The gga-miR_HR_ genes are central to several developmental signalling networks interfacing at Wnt cascade. Amongst gga-miR_HR_ genes several NPS from the Wnt cascade nearly achieved structural features of functional miRNAs. **b,** Application of miR-4673 and activation of the identified gga-miR_HR_ shadow interactome boosted angiogenic activity in chorio-allantoic membrane (top). Blue pseudo-colorized elements demonstrate expansion of microvessels (black arrow) along with increased hematopoiesis. The miRNA also amplified hematopoiesis (red pseudo-colour) and neurogenesis (black arrows) in the explanted embryo (frontal cortex: white arrow) (** indicates p>0.01).

## Discussion

Our findings provide evidence for the evolutionary trajectory of miR-4673 by co-option (as opposed to *de novo* evolution) of genomic elements that operate to reduce transcriptional noise in the Wnt pathway. The proposed model suggests two major steps for the evolution of miR-4673. In the first stage and in the absence of the mature miRNA, selection of intragenic enhancers in Wnt cascade modulator genes seeded homologous primary sequences in a functional cluster of related genes. Throughout the next stage, competition for the reduction of transcriptional noise in Wnt cascade modulator genes (enrichment of TFAP2-binding motifs and palindromic symmetry) propelled structural maturation of miR-4673 in notch-1 while the other competing enhancers partially completed the journey.

Our findings regarding the evolution of pre-miR-4673 genomic sequence from a nucleosome positioning sequence are in agreement demonstrated enrichment of a positioned nucleosome on pre-miRNA genomic sequences (Ozsolak et al., 2008; Zhu et al., 2011). Also in the light of the evidence for co-evolution of miRNA-target pairs (Liu et al., 2016), it is surprising that the targets of miR-4673 demonstrate similar nucleosome positioning features. The co-evolution of miRNA-target pairs from homologous genomic structures predicts a degree of sequence homology that exceeds the reported 6-mer seed region criterion in animals. In a drastic contrast, plant miRNAs often demonstrate near-perfect complementarity to the targeted mRNAs (Jones-Rhoades and Bartel, 2004; Rhoades et al., 2002). While evidence is emerging regarding the importance of miRNA base-pairing beyond the conventional seed region in animal cells (Broughton et al., 2016; Hutvagner and Zamore, 2002), we believe that the dichotomy (seed region length) may in part result from the dynamic evolutionary landscape of plants compared to animals. Unlike animals, plants are stationary and hence rely on constant evolutionary adaptations by genomic innovations (Traverse, 2009). The rapid evolutionary rate of plant genes could provide a more accurate overview of miRNA-target homology for several reasons. The newly evolving microRNAs may exhibit near-perfect homology to the targets due to insufficient time required for mutational drift of the homology region. On the other hand, the redundant drifting miRNA-target interactions that generate the illusion of “seed region” will be eliminated due to the rapid turnover rate of the genetic pool. This line of reasoning fits our observation of the near-perfect complementarity between miR-4673 and some of the targets that leads to the degradation of the targeted transcript, e.g. cdk-18(Dokumcu et al., 2018). We suggest that the recent evolutionary history of miR-4673 leaves insufficient time for mutational drift of the targets thereby replicating the hybridization dynamics of plant microRNAs. In this scenario, the conventional 6-mer-based interactions are less important and belong to the drifting targets under instruction from other genomic forces. The latter hypothesis remains to be validated.

## Conclusions

The findings reported herein attest to a remarkable potential for accelerated reprogramming of genomic connectivity through cis-regulatory elements concealed in non-coding DNA. Major evolutionary adaptations may be generated by subtle modulation of the transcriptional temporal domains (enhancers) that propel evolution of microRNAs. Findings of the present study may have broader implications for understanding the role of non-coding DNA in the evolution of complex metazoans.

## Author’s Contributions

R.M.F. conceived, designed, and performed the experiments, analysed and interpreted data, and wrote the manuscript. S.R.L. performed the experiments. N.H. conceived experiments, contributed to interpretation of the data and contributed to writing the manuscript.

